# Reduced benzoxazinoid defences favour maize beneficial colonisation by *Colletotrichum tofieldiae*

**DOI:** 10.64898/2026.07.31.742045

**Authors:** Carlos González-Sanz, Bo Wang, Sara González-Bodí, Víctor Flors, Laura Rodríguez-Casillas, Sandra Díaz-González, Soledad Sacristán

## Abstract

- Specialised defence metabolites mediate plant–microbe interactions, but their regulation during beneficial associations in cereals remains poorly understood. Here we investigate how maize (*Zea mays*) chemical defences are modulated during interaction with the beneficial fungal endophyte *Colletotrichum tofieldiae* (strain Ct0861) to promote host growth.
- To elucidate the molecular basis of this interaction, we combined transcriptomics, metabolite analyses and functional assays. To assess benzoxazinoids (BXs) roles at the plant–fungus interface, we compared Ct0861 colonisation and growth-promotion in wild-type maize and the BX-deficient *bx1::DS* mutant, using the pathogen *Colletotrichum graminicola* (CgM1.001) as reference.
- Transcriptomic analyses revealed coordinated regulation of specialised defence pathways during the initial stages of Ct0861 colonisation, featuring systemic induction of terpenoid phytoalexin biosynthesis alongside localised suppression of roots BXs. Metabolomic analyses showed reduced apoplastic MBOA in Ct0861-colonised roots. Toxicity assays evidenced that Ct0861, unlike CgM1.001, is highly sensitive to MBOA. Accordingly, Ct0861 accumulation and growth-promoting effects were significantly enhanced in the *bx1::DS* mutant without causing disease symptoms.
- Our findings indicate that BXs restrict Ct0861 maize colonisation, where BX deficiency increases fungal biomass and enhances growth promotion. Understanding how crops fine tune specialised metabolism to balance microbial restriction with beneficial accommodation provides valuable insights for sustainable agriculture.

## INTRODUCTION

Crops are constantly exposed to a diverse community of organisms inhabiting both belowground and aboveground compartments (Díaz-González *et al*., 2025a). To cope with this complexity, plants have evolved sophisticated chemical defence systems that not only protect against pathogens and herbivores but also mediate interactions with beneficial microbes (Erb & Kliebenstein, 2020; Jacoby *et al*., 2021). These chemically mediated interactions are highly dynamic and context-dependent, often determining whether an encounter results in disease or mutualism (Hacquard *et al*., 2017). While the role of these metabolites in restricting pathogen invasion has been well characterised, their regulation during beneficial plant–fungus interactions remain largely unexplored, particularly in cereal crops such as maize (*Zea mays*).

In maize, chemical defence against microbial invaders relies on three major classes of specialised metabolites: terpenoid phytoalexins, phenylpropanoid-derived compounds and benzoxazinoids, each with distinct biosynthetic origins and ecological roles (Yasmin *et al*., 2024). Maize produces a diverse terpenoid arsenal through the coordinated activity of multiple terpene synthases (TPSs) and cytochrome P450 monooxygenases (CYPs), which collectively generate structurally diverse antimicrobial metabolites, including sesquiterpenoid and diterpenoid phytoalexins such as zealexins (ZXs), kauralexins (KXs), and dolabralexins (DXs) (Block *et al*., 2019). These compounds are rapidly induced upon pathogen attack and exhibit strong antifungal activity (Schmelz *et al*., 2011; Ding *et al*., 2020). Beyond their defensive roles, maize terpenoids have been associated with drought tolerance (Vaughan *et al*., 2015) and may also contribute to root development (Murphy *et al*., 2023). In addition to terpenoids, maize produces a subset of flavonoids, a large group of phenylpropanoid and polyketide-derived metabolites, which also accumulate following pathogen infection and contribute to disease resistance (Förster *et al*., 2022).

Another major class of maize metabolites comprises the benzoxazinoids (BXs), specialised indole-derived defence metabolites produced predominantly by grasses (Poaceae) (Niculaes *et al*., 2018). BXs fulfil diverse ecological functions, ranging from defence against herbivores and pathogens to roles in rhizosphere interactions and allelopathy (Zhou *et al*., 2018). These compounds are constitutively accumulated in young seedlings and typically decline during plant development. BX biosynthesis starts from indole-3-glycerol phosphate (IGP), a central precursor in maize indole metabolism that is also connected to tryptophan and indole-3-acetic acid (IAA) biosynthesis. In the BX pathway, IGP is converted into indole by the chloroplast-localised BX1 enzyme (Richter *et al*., 2021). Indole is subsequently modified through a series of reactions involving CYP enzymes to generate intermediates such as 2,4-dihydroxy-1,4-benzoxazin-3-one (DIBOA), which are then glycosylated in the cytoplasm, allowing their storage in a stable and inactive form (Robert & Mateo, 2022). BX glucosides are stored in the vacuole until cell disruption, after which aglycones are released (Niculaes *et al*., 2018). Upon tissue disruption, BX glucosides are hydrolysed by β-glucosidases, releasing unstable aglycones such as DIBOA and 2,4-dihydroxy-7-methoxy-1,4-benzoxazin-3-one (DIMBOA). These compounds can spontaneously degrade into the corresponding benzoxazolinones: DIBOA mainly gives rise to 1,3-benzoxazol-2(3H)-one (BOA), whereas DIMBOA is converted into the more stable 6-methoxy-benzoxazolin-2-one (MBOA). BX biosynthesis is highly compartmentalized within the cell, with early steps occurring in the chloroplast, intermediate reactions in the endoplasmic reticulum, and later hydroxylation and glycosylation steps taking place in the cytosol and chloroplasts (Zhao *et al*., 2024).

Besides their intracellular synthesis, BXs and related metabolites also accumulate and are metabolized in the apoplast, especially under biotic stress. Pathogen-associated elicitors, such as chitosan, enhance the accumulation of DIMBOA and 2-hydroxy-4,7-dimethoxy-1,4-benzoxazin-3-one glucoside (HDMBOA-Glc) in the apoplast without wounding, indicating an active and regulated export of BXs into the extracellular space (Ahmad *et al*., 2011). In this context, apoplastic DIMBOA can act as a defence-related signal by promoting callose deposition, thereby strengthening penetration resistance against aphids and fungal pathogens. Beyond the apoplast, BXs can be released into the rhizosphere through root exudation (Hassan & Mathesius, 2012; Murphy *et al*., 2021; Gfeller *et al*., 2023), where they act as antimicrobial and signalling molecules that influence the assembly, composition, and activity of root-associated microbiota (Kudjordjie *et al*., 2019; Cotton *et al*., 2019; Cadot *et al*., 2021; Murphy *et al*., 2021). This selection process favours BX-tolerant or BX-detoxifying endophytes (Saunders & Kohn, 2009; Neal *et al*., 2012; Schütz *et al*., 2019; Thoenen *et al*., 2023). For instance, the BX breakdown product MBOA shows strong fungistatic activity against multiple cereal pathogens, including *Fusarium* spp.; however, several maize-associated *Fusarium* species display high tolerance and the ability to detoxify BOA/MBOA, which may contribute to their frequent symptomless endophytic lifestyle (Glenn *et al*., 2001; Saunders & Kohn, 2009). Thus, maize chemical defences not only restrict pathogen proliferation but also define microbial niches available for beneficial root-associated microorganisms.

Among beneficial fungal endophytes, *Colletotrichum tofieldiae* has emerged as a model system to study the integration of plant nutrition and immunity in plant-endophyte interactions. The strain Ct0861 was originally isolated from surface-sterilised leaves of asymptomatic *Arabidopsis thaliana* plants collected from a natural population in central Spain (García *et al*., 2013). Previous work demonstrated that Ct0861 promotes plant growth in *A. thaliana* by transferring phosphorus to the host in a phosphorus-dependent manner (Hiruma *et al*., 2016). This growth-promoting effect depends on the integration of the plant phosphate starvation response (PSR) with the indole glucosinolate (IG) biosynthetic pathway (Hiruma *et al*., 2016). IGs are tryptophan (Trp)-derived defence metabolites exclusive to Brassicaceae that play a central role in modulating Ct0861 colonisation. This is evidenced by the shift from beneficial to pathogenic behaviour observed in the mutant *cyp79b2b3*, which is impaired in the biosynthesis of these compounds (Hiruma *et al*., 2016).

However, the host range of Ct0861 extends beyond Brassicaceae, as it colonises and promotes growth in crop species such as tomato and maize (Díaz-González *et al*., 2020). Field trials have shown that Ct0861 significantly enhances maize yield under sufficient phosphate conditions. Moreover, maize plants colonised by Ct0861 show reduced biomass of the mycotoxigenic fungus *Aspergillus flavus* in the cobs, suggesting the induction of systemic defences (Díaz-González *et al*., 2025b). Nonetheless, despite its demonstrated agronomic relevance, it remains unknown how Ct0861 interacts with maize specialised defence pathways to achieve colonisation and promote host growth.

Here, we investigate the transcriptional and metabolic reprogramming of maize during early colonisation by Ct0861. By integrating transcriptomic analyses across tissues and time points with targeted functional assays, we examine how maize chemical defence pathways are remodelled during initial endophyte establishment and to what extent these responses differ from those triggered by fungal pathogens. Given their role as major indole-derived metabolites in grasses, we evaluated whether BXs contribute to the regulation of Ct0861 colonisation and its beneficial effects in maize. Our results suggest a coordinated transcriptional regulation between indolic and terpenoid defences during the onset of this beneficial association, pointing to balancing roles that may shape the outcome of plant–endophyte interactions.

## MATERIALS AND METHODS

### Plant and fungal materials

The maize (*Zea mays* L.) variety LG 34.90 (Limagrain Ibérica, SA, Pamplona, Spain) was used as the reference for all the experiments unless otherwise specified. The *BX1* gene knockout mutant *bx1::DS* (W22-T43) and its near-isogenic line W22 (T43) (Tzin *et al*., 2015; Hu *et al*., 2018) were kindly provided by Dr. Georg Jander (Boyce Thompson Institute, NY, USA).

*Colletotrichum tofieldiae* (Pat.) Damm, P.F. Cannon & Crous strain Ct0861 was isolated from natural *Arabidopsis thaliana* (L.) Heynh. populations in the central Iberian Peninsula (García *et al*., 2013). *Colletotrichum graminicola* (Ces.) G.W. Wilson M1.001 strain was provided by Dr. Serenella Sukno from University of Salamanca (Spain) (Sukno *et al*., 2008).

### Growth conditions and fungal inoculations

To prepare Ct0861 inoculum, PDA plates (Difco^TM^ Becton, Dickinson and Co., Maryland, USA) were inoculated at the centre with a 3 µL droplet of a conidial suspension (10^6^ conidia/mL) obtained from a 10^8^ conidia/mL stock stored at −80°C in 2% skimmed milk (Sveltesse®, Nestlé España, Barcelona, Spain). To prepare CgM1.001 inoculum, PDA plates (Difco^TM^ Becton, Dickinson and Co., Maryland, USA) were inoculated at the centre with a 3 µL droplet of a conidial suspension (10^6^ conidia/mL) stock stored at −80°C in 30% glycerol. Inoculated plates were incubated in a growth chamber at 24°C under a light cycle of 14 h light/10 h darkness for Ct0861 and at 28°C in continuous darkness for CgM1.001.

For *in vitro* growth, maize seeds were sterilised by rinsing them for 10 min in 70% ethanol and 15 min in 20% bleach, followed by three washes of 3 minutes in sterile deionized water and then dried. To achieve the desired inoculum dose of 10^3^ conidia per seed, 700 μL of a conidial suspension (1.4 × 10⁵ conidia/mL) or water (for controls) was added to 100 seeds in 50 mL Falcon tubes and vigorously shaken to ensure uniform distribution on the seed surface. Subsequently, five seeds were placed in square Petri plates containing ½ MS medium (Caisson Laboratories) supplemented with Microagar (Duchefa Biochemie), adjusted to pH 5.7 and incubated in complete darkness at 24°C.

For metabolomic measurements of roots and apoplastic fluid, maize seeds were inoculated as described above and placed in pots with vermiculite (one seed per pot). Plants were watered once per week with 100 mL of Hoagland’s solution and once per week with sterile deionized water. Plants were cultivated in a growth chamber (HERAEUS, VB 1514, Vötsch Industrietechnik GmbH, Hanau, Germany) at 24-21°C (light/dark), with long-day cycle of 14 h light, 140 μmol m⁻² s⁻¹and 65% RH. After 7, 11 and 15 days after sowing, roots from 5 plants were cleaned carefully, collected and directly frozen in liquid nitrogen or used for apoplastic fluid isolation as described below.

For RNA extraction of axenic fungal cultures, Ct0861 and CgM1.001 were grown on sterile cellophane membranes placed on the surface of ½ MS agar plates, using the same medium as that used for maize seedling growth. Five 5 µL droplets of a 10⁴ conidia/mL conidial suspension were deposited onto each membrane. Plates were incubated under the same conditions described above for maize experiments. After 7 days, the membrane was carefully collected for RNA extraction.

### RNA extraction, sequencing and differential expression analysis

For RNA-seq analysis, plants grown *in vitro* were harvested at 4 and 7 dpi. Roots and shoots from 5 plants were pooled per biological replicate (3 biological replicates per condition) and immediately flash-frozen in liquid nitrogen. RNA was extracted with ROTI®-phenol (Carl Roth, Karlsruhe, Germany) and residual DNA was removed by DNase I (Thermo Fisher Scientific, Schwerte, Germany) according to the manufacturer’s instructions.

RNA samples were sent for mRNA-Seq analysis to Novogene UK CL (Cambridge, UK). Poly(A)-selected mRNA libraries were prepared and sequenced with Illumina technology (2×150 bp paired-end reads), generating over 50 million read pairs per sample.

STAR software v2.7.11b (Dobin *et al*., 2013) was used to align the reads to the *Z. mays* reference genome (Zm-B73-REFERENCE-NAM-5.0). Raw read counts per gene were obtained using HTSeq v2.0.8 (Anders *et al*., 2015). Downstream analyses were performed using RStudio software (v4.2.2) and R (v4.3.2). Differential expression analysis was performed with Bioconductor (v3.20) package DESeq2 v1.46.0 (Love *et al*., 2014). Multiple testing correction was applied using the Benjamini–Hochberg (BH) procedure, as implemented in DESeq2. Significantly differentially expressed genes (DEGs) were considered those with a |log2FC| > 1 and *padj* < 0.05. Venn diagrams were generated using the R package VennDiagram v1.7.3. Heatmaps were generated using the R package ComplexHeatmap v2.22.0.

Maize genes were mapped to Gene Ontology (GO) and Kyoto Encyclopedia of Genes and Genomes (KEGG) using three databases: AgriGO v2.0 (Tian *et al*., 2017), Ensembl Plants (Dyer *et al*., 2025) release 59 and InterProScan (Jones *et al*., 2014). The package clusterProfiler v4.14.4 was used to perform GO and KEGG enrichment analysis on the DEGs. GO and KEGG terms were considered overrepresented if their false discovery rate (FDR) was ≤ 0.05. Redundant GO terms were removed using REVIGO (Supek *et al*., 2011). Enrichment plots were generated using the R package ggplot2 v3.5.2.

### Quantitative RT-PCR analysis (qRT-PCR)

Roots and shoots from five plants grown *in vitro* were collected and frozen in liquid nitrogen immediately. For axenic fungal samples, the cellophane membranes were carefully removed and immediately frozen in liquid nitrogen. All samples were ground to a fine powder using a mortar and pestle in liquid nitrogen, and RNA was extracted as described above. One µg of total RNA was used as template for cDNA synthesis using the Transcriptor First Strand cDNA Synthesis Kit (Roche Diagnostics GmbH, Mannheim, Germany). qRT-PCR reactions were performed using LightCycler 480 SYBR Green I Master (Roche Diagnostics GmbH, Mannheim, Germany) and gene-specific primers listed in Table S1. For maize gene expression analysis, *ZmACTIN* was used as the reference gene. For fungal gene expression analysis, *CtBETA-TUBULIN* for Ct0861 (Hiruma *et al*., 2016) and *CgH3* (Krijger *et al*., 2008) for CgM1.001 were used for normalization.

### Apoplastic fluid isolation

Apoplastic fluid from maize roots at 7, 11 and 15 dpi of plants grown in pots as described above was extracted following a previously described protocol with minor modifications (Yu *et al*., 1999). Briefly, roots were collected and carefully washed with Milli-Q water to remove attached vermiculite and dried with paper towels. Roots from 8-12 plants were pooled and immediately weighed. The primary roots were then cut into 2 cm segments, placed in sterile ice-cold Milli-Q water, and subjected to two rounds of 15 min of vacuum infiltration. To collect apoplastic fluid, root samples were wrapped in Parafilm^TM^ (Amcor Flexibles Northamerica, Wisconsin, USA), placed into 10 mL syringes inserted into 50 mL Falcon tubes, and centrifuged for 15 min at 200 *x g* at 4 °C. The flowthrough was frozen and subsequently freeze-dried. To assess cytoplasmic contamination, an aliquot of each sample was analysed using an MDH assay, as described in Rodríguez de Lope et al. (2025).

### Benzoxazinoid extraction and quantification

For BX extraction, previously described protocols were followed with minor modifications (Sasai *et al*., 2009; Ahmad *et al*., 2011). Briefly, 50 mg aliquots of lyophilized plant material were weighed into microcentrifuge tubes. Metabolites were extracted by adding 1 mL of methanol acid water (49:1:50, v/v/v) supplemented with N-benzoyl-L-tyrosine (10 µg/L) and d₂-IAA as internal standards. After buffer addition, the samples were homogenised under cold conditions using a TissueLyser system (Qiagen, Hilden, Germany) for 30 s and subsequently incubated on ice for 10 min to ensure complete tissue hydration. The homogenate was then centrifuged at full speed for 10 min. The resulting supernatant was collected, filtered through a 0.22 µm nylon membrane filter, and transferred into a clean UPLC vial prior to chromatographic analysis. A 5-µL aliquot of a 1:3 dilution was injected into an Acquity UPLC system (Waters, Milford, MA, USA) coupled with a triple quadrupole mass spectrometer using a UPLC Kinetex 2.6 μm EVO C18 100 A, 2.1 × 50 mm (Phenomenex) column. External calibration curves were prepared with pure standards of IAA, HDMBOA-Glc, MBOA, DIMBOA and DIMBOA-Glc. All BXs standards were kindly provided by Prof Ton (Sheffield University, UK) and Prof Roberts (Bern University, Switzerland).

The transitions used were selected in ESI+ mode as follows: 176.1>130.1 for IAA; 388>226 for HDMBOA-Glc; and in ESI-mode as follows: 164>149 for MBOA; 210> 164 for DIMBOA and 372>210 for DIMBOA-Glc. The chromatographic separation of BXs was achieved using a binary mobile phase system consisting of Mobile Phase A (100% Milli-Q water) and Mobile Phase B (methanol:isopropanol:acetic acid, 3800:200:1, v/v/v).

### MBOA toxicity assays

BXs tolerance assays were performed as described by Ridenour & Bluhm (2017), with slight modifications. Stock solutions of MBOA (Thermo Fisher Scientific, Schwerte, Germany) were prepared with dimethyl sulfoxide (DMSO). Four-millimetre agar plugs from 7-day-old Ct0861 or CgM1.001 cultures were placed at the centre of PDA plates amended with corresponding concentrations of DMSO as controls and different concentrations of MBOA. Plates were incubated at 24 °C under (light/dark conditions). Growth inhibition was calculated relative to the corresponding DMSO control growth after 10 days.

The amino acid sequence of *GLRG_10902* from *C. graminicola* was used as a query in a DIAMOND v2.2.1 (Buchfink *et al*., 2014) blastp search against the predicted Ct0861 proteome (Hiruma *et al*., 2023). *Ct61P_02057* was selected as the closest putative homolog based on sequence similarity, query coverage, and E-value. Nucleotide sequences of both *GLRG_10902* and *Ct61P_02057* were used to design primers (Table S1) to analyse their expression by qRT-PCR. For that, four-mm agar plugs from 7-day-old Ct0861 or CgM1.001 cultures were transferred to 24-well plates containing 1 mL of sterile liquid ½ MS supplemented with 0.25 and 0.5 mg/mL concentration of MBOA. Control wells contained the corresponding concentration of DMSO without MBOA. Plates were incubated at 24°C in darkness for 48 h. After incubation, fungal mycelia were collected and immediately frozen in liquid nitrogen. This material was used to analyse the expression of the selected candidate MBOA-detoxification genes by qRT-PCR as described above.

### Colonisation assays

Maize seeds were inoculated and grown *in vitro* as described above. After 7 days, roots from 5 plants from each plate were cleaned thoroughly with tap water, dried with sterile filter paper and frozen in liquid nitrogen. Tissues were ground with mortar and pestle, DNA was extracted using CTAB method (Murray & Thompson, 1980). Fungal biomass was quantified by qPCR using specific primers for Ct0861 or for CgM1.001 (**Table S1**). Samples were standardised against *ZmBETA-TUBULIN* gene (Mideros *et al*., 2009).

### Statistical analysis

Statistical analyses were performed in RStudio software v4.2.2. Data distribution and homogeneity of variances were assessed using the Shapiro–Wilk test and Bartlett’s test, respectively. When data met the assumptions of normality and homoscedasticity, differences between two groups were analysed using Student’s *t*-test, whereas comparisons among more than two groups were performed using one-way analysis of variance (ANOVA), followed by Fisher’s least significant difference (LSD) post hoc test with Bonferroni correction for multiple comparisons. When data did not meet the assumptions of normality, differences between two groups were analysed using the Wilcoxon rank-sum test, whereas comparisons among more than two groups were performed using the Kruskal–Wallis test, followed by Dunn’s post hoc test with Bonferroni correction for multiple comparisons. Differences were considered statistically significant at *p* < 0.05. Permutational multivariate analyses of variance (PERMANOVA) were performed using the ‘adonis2’ function implemented in the vegan package (version 2.6.4). Graphs were generated using the ggplot2 package (v3.5.2).

## RESULTS

### Early transcriptional response of maize to Ct0861 colonisation

To investigate the early transcriptional response of maize to colonisation by Ct0861, RNA-seq analyses were performed. First, we conducted a time-course experiment measuring root and shoot length in plants grown *in vitro* to select the optimal time points for our analysis (**Fig. 1a**). Four dpi was selected as the first time point, since it preceded the observation of growth promotion effects, whereas 7 dpi was chosen as the second time point when significant differences in root and shoot length were observed compared to the mock treatment.

**Figure 1.**
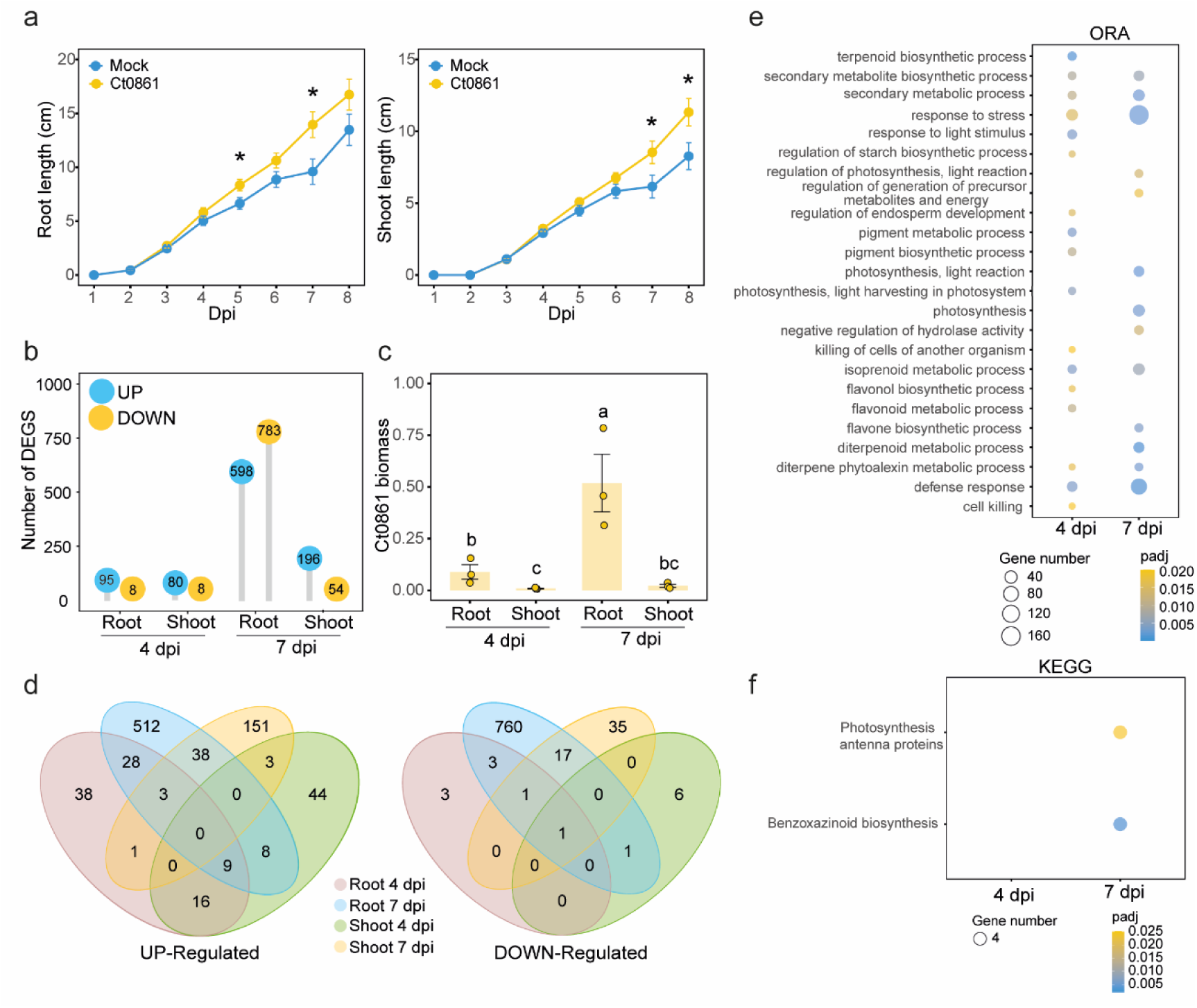
Comparative analysis of the transcriptomic response in maize (*Zea mays*) roots and shoots with *Colletotrichum tofieldiae* strain Ct0861 seed inoculation after 4 and 7 days post inoculation. **a)** Time course of root (left) and shoot (right) length of maize seedlings after seed inoculation with Ct0861 spores. Statistical significance was assessed Student’s t-test. Asterisks indicate statistically significant differences (P < 0.05). **b)** Lollipop plot showing the number of differentially expressed genes (DEGs) (|log2FC| > 1 and *padj* < 0.05) in maize tissues (roots and shoots) upon Ct0861 seed treatment compared with the mock treated plants at two different time points (4 and 7 dpi). **c)** Ct0861 biomass determination in the different samples by qPCR using a fungus-specific primer relative to a maize reference gene (β-tubulin). Values represent the mean ± standard error of 3 biological replicates. **d)** Venn diagrams showing the overlap between the up regulated and down regulated DEGs between the different samples. **e)** Gene Ontology (GO) biological process enrichment analysis of DEGs in maize roots at 4 and 7 dpi. **f)** Kyoto Encyclopaedia of Genes and Genomes (KEGG) pathway enrichment analysis of DEGs identified in maize roots at 4 and 7 dpi.

Using these selected time points, we generated RNA-seq datasets from shoot and root samples of plants grown with or without Ct0861. To explore the overall plant transcriptomic response, principal component analysis (PCA) was performed using normalized expression (**Fig. S1a**). The samples clustered primarily by tissue type along the horizontal axis, which explained 81% of the variance, and secondarily by time point along the vertical axis, which explained 8% of the variance. PERMANOVA supported this clustering pattern, identifying tissue type as the main factor explaining global transcriptomic variation (R² = 0.808, *P* < 0.001), followed by time point (R² = 0.068, *P* < 0.001). The heatmap with the normalised counts for all the libraries also showed the main clustering due to tissue and time of sampling, but also revealed substantial transcriptomic reprogramming triggered by fungal interaction, especially in root samples at 7 dpi, where mock and Ct0861 inoculated samples did not cluster together (**Fig. S1b**).

To decipher the molecular changes induced by Ct0861 in maize tissues, differential expression analyses were conducted comparing data from Ct0861 inoculated plant samples to those from mock-treated controls at each time point and tissue type (**Table S2**). At 4 dpi, we observed a slight transcriptional response, with 103 and 88 DEGs identified in roots and shoots, respectively (**Fig. 1b**). In contrast, at 7 dpi, a more robust response was detected in roots, with 1379 DEGs (596 upregulated and 783 downregulated). Shoots also showed an increase in the number of DEGs at 7 dpi compared to 4 dpi, with a total of 250 DEGs. To estimate fungal biomass in each sample, we quantified it by qPCR using specific primers for Ct0861 (gene *CT0861_00602*). Results indicated a progressive increase in fungal biomass in roots over time, whereas fungal biomass in shoots remained stable (**Fig. 1c**). Venn diagrams identified shared or unique upregulated and downregulated DEGs across time (4 or 7 dpi) or plant tissue (root or shoot) (**Fig. 1d**). The analysis revealed a highly tissue- and time-specific transcriptional response, being most of DEGs specific of each type of sample. Notably, no common upregulated DEGs were found across all four conditions, while only one downregulated DEG was shared.

To gain further insight into the biological processes and pathways associated with transcriptional responses, we performed GO and KEGG enrichment analyses on the DEGs list (**Table S3**). At the earlier time point (4 dpi) in roots, we identified enriched GO categories predominantly associated with plant defence, including response to stress (GO:0006950), defence response (GO:0006952), and diterpene phytoalexin metabolic process (GO:0051501). Additionally, more specific categories related to programmed cell death were also enriched. Several categories were associated with the metabolism of secondary metabolites, including secondary metabolic process (GO:0019748), terpenoid biosynthetic process (GO:0016114), isoprenoid metabolic process (GO:0006720), and flavonoid biosynthetic process (GO:0009813), all of which are linked to defence pathways. Several categories related to photosynthesis were also overrepresented, such as photosynthesis (GO:0015979), pigment biosynthetic process (GO:0046148), and pigment metabolic process (GO:0042440). At this time point, only one KEGG pathway was significantly enriched in shoots: biosynthesis of secondary metabolites (KEGG:01110). These findings highlight the early recognition of fungal presence and the activation of plant defence mechanisms.

At 7 dpi, roots displayed the highest number of enriched categories, correlating with the increased number of DEGs (**Fig. 1e; Table S3**). The enriched categories were largely similar to those observed at 4 dpi, but containing greater numbers of genes. There were GO categories related to the biosynthesis of secondary metabolites involved in defence, such as secondary metabolite biosynthetic process (GO:0044550), diterpenoid biosynthetic process (GO:0016102), phytoalexin metabolic process (GO:0052314), flavone biosynthetic process (GO:0051553), and isoprenoid metabolic process (GO:0006720). Similar to 4 dpi, we identified categories related to photosynthesis, including photosynthesis, light reaction (GO:0019684), photosynthesis (GO:0015979), and regulation of photosynthesis, light reaction (GO:1901896). Additionally, some enriched categories were found at 7 dpi, including lipid biosynthetic process (GO:0008610), response to karrikin (GO:0080167), negative regulation of proteolysis (GO:0045861), and negative regulation of hydrolase activity (GO:0051346). KEGG enrichment analysis revealed only two enriched pathways: photosynthesis antenna proteins (KEGG:00196) and benzoxazinoid biosynthesis (KEGG:00402) (**Fig. 1f**). These results indicate a strong transcriptomic modulation of genes involved in plant secondary metabolism.

In shoots, the transcriptomic response at 7 dpi was predominantly associated with detoxification of reactive oxygen species (ROS). Enriched GO categories included reactive oxygen species metabolic process (GO:0072593), hydrogen peroxide catabolic process (GO:0042744), response to oxidative stress (GO:0006979), cellular oxidant detoxification (GO:0098869), and response to toxic substances (GO:0009636). Additionally, we identified categories related to cell wall organization and biogenesis (GO:0071554), as well as general metabolic processes such as cellular catabolic process (GO:0044248), transmembrane transport (GO:0055085), secondary metabolic process (GO:0019748), and secondary metabolite biosynthetic process (GO:0044550). KEGG enrichment analysis revealed pathways related to phenylpropanoid biosynthesis (KEGG:00940), biosynthesis of secondary metabolites (KEGG:01110), and metabolic pathways (KEGG:01100).

Collectively, these results emphasize the transcriptional reprogramming in maize tissues in response to Ct0861, with a stronger and more extensive modulation observed in roots at 7 dpi, especially related to secondary metabolism biosynthetic pathways of defence related molecules such as terpenoids, BXs, and phenylpropanoids.

### Ct0861 modulates chemical defence pathways in maize roots

We generated heatmaps from the transcriptomic data to display the genes expressed in the root involved in various maize defence metabolic pathways: DXs and KXs (diterpenoids), ZXs (sesquiterpenoids) and BXs (indolic compounds) (**Fig. 2a**). KXs and DXs genes showed modest upregulation at 4 dpi, and stronger induction at 7 dpi. ZX biosynthetic genes followed a similar pattern to that of KXs and DXs, although their upregulation was more pronounced after 4 days of interaction. However, the BXs heatmap shows downregulation in several genes only at 7 dpi.

**Figure 2.**
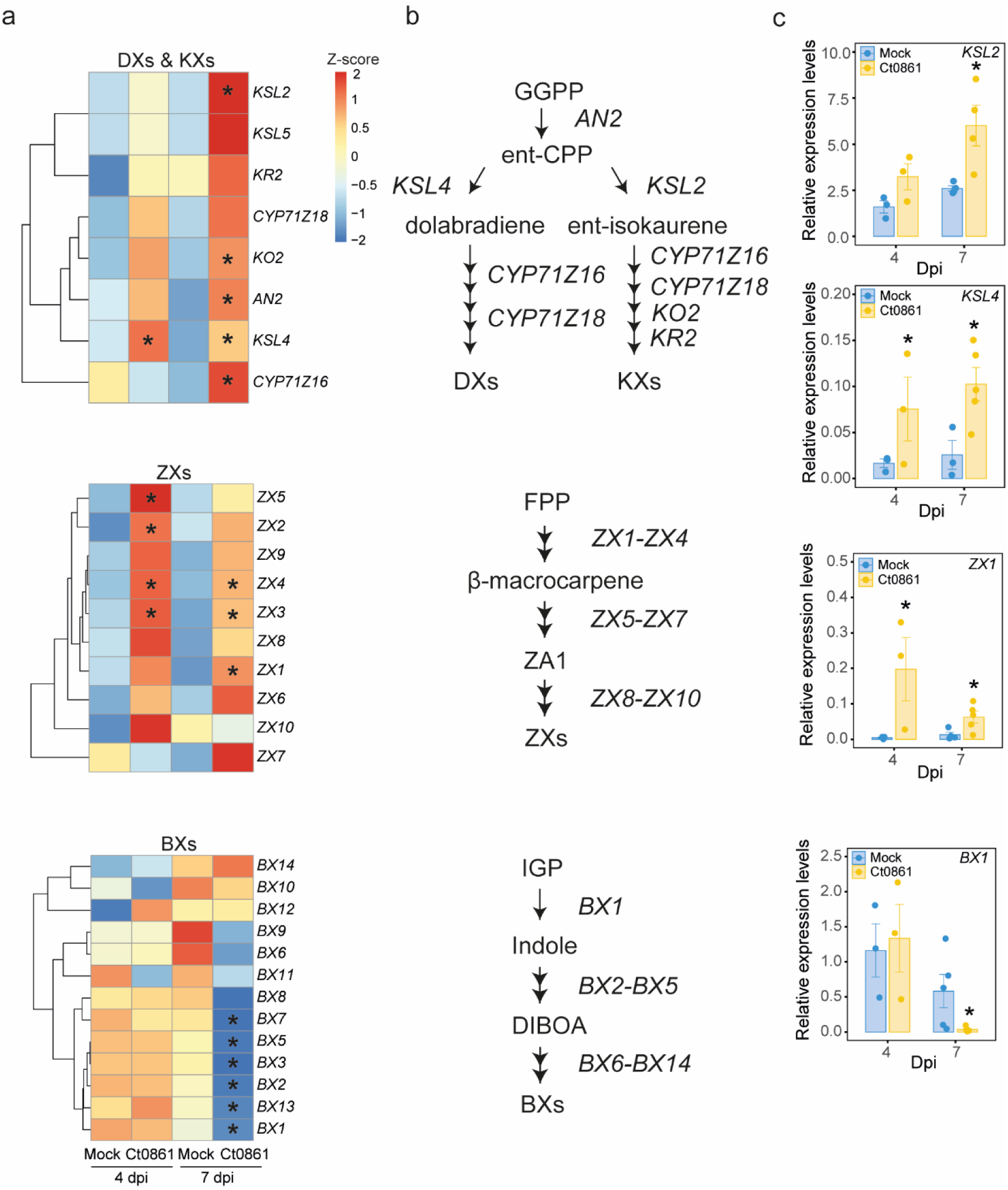
*Colletotrichum tofieldiae* strain Ct0861 modulates the expression of maize (*Zea mays*) chemical defence biosynthetic pathways. **a)** Heatmaps showing RNA-seq expression patterns of genes involved in the biosynthesis of benzoxazinoids (BXs), dolabralexins (DXs), kauralexins (KXs), and zealexins (ZXs) in roots of mock- and Ct0861-inoculated maize seedlings at 4 and 7 days post-inoculation (dpi). Gene expression values are represented as row-scaled Z-scores. Asterisks indicate significant differences between Ct0861-inoculated and mock-treated roots at the corresponding time point, as determined by a likelihood ratio test followed by Benjamini–Hochberg correction (*padj* < 0.05). **b)** Simplified representation of the BX, DX, KX, and ZX biosynthetic pathways in maize. **c)** Relative expression levels of selected genes involved in the early steps of the DX/KX, ZX, and BX biosynthetic pathways, quantified by RT–qPCR using gene-specific primers. *ZmACTIN* was used as the reference gene. Bars represent the mean ± standard error. Internal line in box-plot boxes indicates the median or second quartile (Q2), upper line of the boxes indicates the third quartile (Q3) and lower line the first quartile (Q1) of the data. Mean is represented by a black dot. Asterisks indicate statistically significant differences in Student’s t-test (*P* < 0.05).

We selected relevant genes involved in the initial steps of each pathway to analyse their expression in maize roots by RT-qPCR in an independent experiment. The genes selected were *KAURENE SYNTHASE-LIKE 4* (*KSL4)* for DXs, *KAURENE SYNTHASE-LIKE 2* (*KSL2)* for KXs (Murphy *et al*., 2023), *TERPENE SYNTHASE 6 (ZX1)* for ZXs (Ding *et al*., 2020) and *BENZOXAZINELESS 1* (*BX1)* for BXs biosynthesis pathways (Ahmad *et al*., 2011). *BX1* and *ZX1* encode the first enzymes in the BXs and ZXs pathways, respectively, whereas *KSL4* and *KSL2* catalyse the branching point leading to the DX and KX pathways from ent-copalyl diphosphate (**Fig. 2b**) (Murphy *et al*., 2023). Results showed that *KSL2* was overexpressed in Ct0861 inoculated roots compared to mock, and this overexpression increased with the time of interaction, being significant after 7 dpi with Ct0861 (**Fig. 2c**). In the case of *KSL4,* upregulation was detected at both 4 dpi and 7 dpi. *ZX1* showed upregulation under colonisation at both time points, with greater differences compared to the mock condition in the earlier time point. Finally, *BX1* exhibited an opposite pattern: no changes were observed at the early time point, but a strong repression was detected at 7 dpi. These results confirm the RNA-seq data, highlighting the distinct temporal regulation of maize antimicrobial defence pathways during interaction with Ct0861 and the specific repression of BX biosynthetic pathway in maize roots at 7 dpi.

### Benzoxazinoid repression occurs without global reprogramming of indole metabolism

Given the specific repression of *BX1*, we investigated whether alternative indole-derived metabolic branches could be activated as a compensatory response. We first analysed the expression of *BENZOXAZINELESS 2* (*BX2*), which acts immediately downstream of *BX1* in the benzoxazinoid biosynthetic pathway and was also downregulated (**Fig. 3a**). We then examined *INDOLE-3-GLYCEROL PHOSPHATE LYASE* (*IGL*) and several *TRYPTOPHAN SYNTHASE (TSA/B)* genes, which can also convert indole-3-glycerol phosphate to indole (Richter *et al*., 2021). *IGL* is important for the synthesis of volatile indole (Ahmad *et al*., 2011), whereas *TSA*, followed by *TSB1, TSB2 and TSB2C* participates in the early steps of tryptophan biosynthesis (Ye *et al*., 2019). RT-qPCR analyses showed no significant changes in the expression of these genes in roots or shoots at either time point (**Fig. 3a; Fig. S2**). Consistently, quantification of indole-3-acetic acid (IAA) levels revealed no differences between Ct0861-colonised and mock-treated plants (**Fig. 3b**). Together, these results indicate that repression of the BX pathway is not associated with a general redirection of indole metabolism.

**Figure 3.**
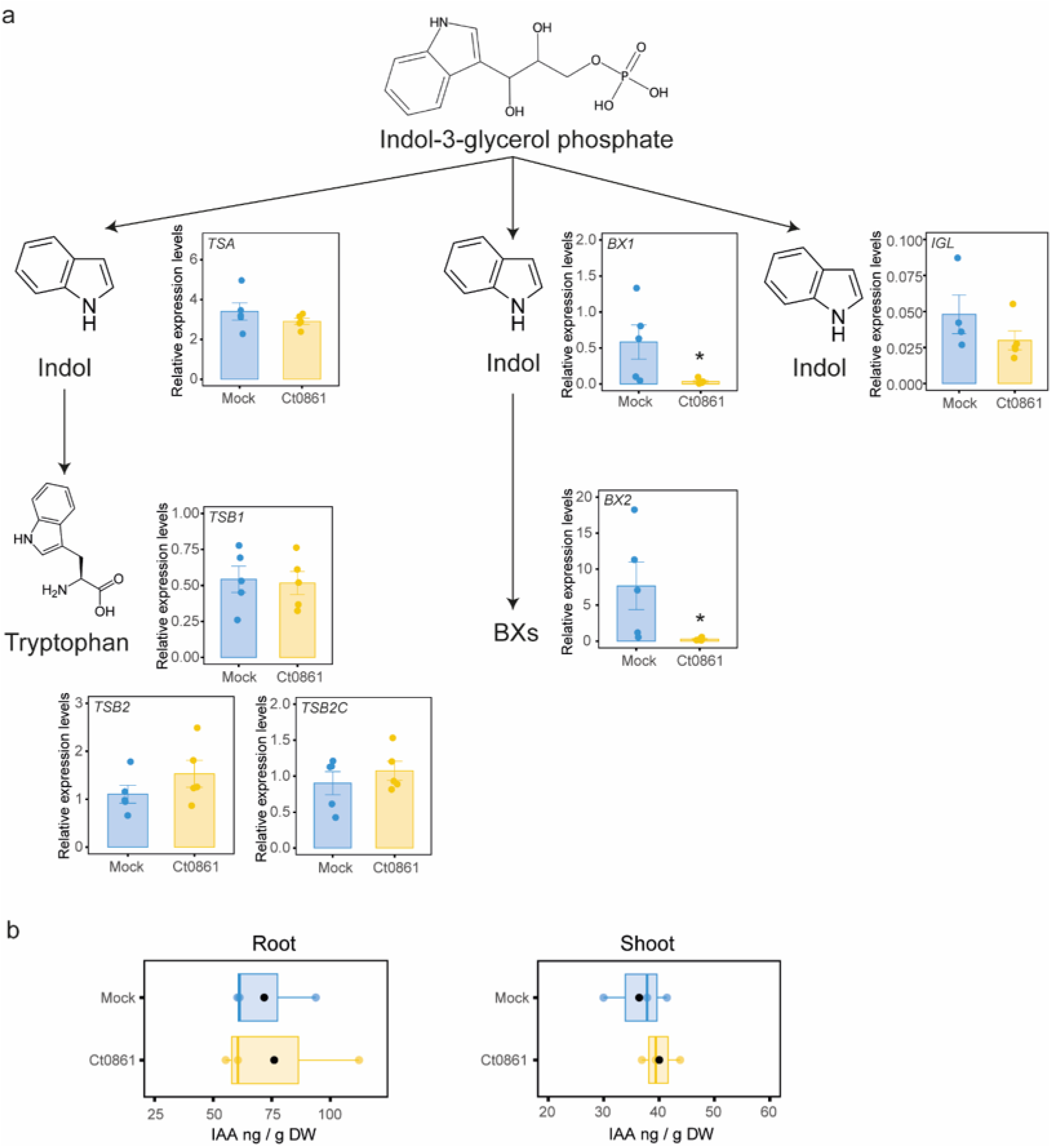
*Colletotrichum tofieldiae* strain Ct0861 affects the expression of genes involved in indole metabolism without altering indole-3-acetic acid (IAA) accumulation in maize (*Zea mays*) seedlings. **a)** Simplified representation of the metabolic routes derived from indole-3-glycerol phosphate leading to tryptophan and benzoxazinoid (BX) biosynthesis. Relative expression levels of *TSA, TSB1, TSB2, TSB2C, BX1, BX2*, and *IGL* in roots of mock- and Ct0861-inoculated maize seedlings at 7 days post-inoculation were quantified by RT–qPCR using gene-specific primers. *ZmACTIN* was used as the reference gene. Bars represent the mean ± standard error. Asterisks indicate significant differences between mock- and Ct0861-inoculated roots (Student’s t-test, *P* < 0.05). **b)** IAA levels in roots and shoots of mock- and Ct0861-inoculated maize seedlings at 7 days post-inoculation measured by UPLC-QToF-MS. Internal line in box-plot boxes indicates the median or second quartile (Q2), upper line of the boxes indicates the third quartile (Q3) and lower line the first quartile (Q1) of the data. Mean is represented by a black dot. No statistically significant differences were detected between treatments (Student’s t-test, *P* > 0.05).

### Ct0861 colonisation transiently reduces apoplastic MBOA levels

We next quantified BX-related compounds in the tissue and apoplastic fluid collected from roots of maize plants grown in pots at different time points during early Ct0861 colonisation (**Fig. 4**). In the roots, the stored glycosylated compounds DIMBOA-Glc and HDMBOA-Glc were the most abundant, but followed opposite trends over time, with DIMBOA-Glc decreasing and HDMBOA-Glc increasing (**Fig. 4a**). The aglycone DIMBOA showed a similar trend to its glycosylated form, whereas its degradation product MBOA subsequently increased. No statistically significant differences were found between Ct0861-inoculated and mock-treated plants. In contrast to whole root tissue, the most abundant compounds in the root apoplastic fluid were the aglycone DIMBOA and its more toxic degradation product, MBOA (**Fig. 4b**). However, while the glycosylated forms (DIMBOA-Glc and HDMBOA-Glc) remained stable at low levels, the high initial levels of DIMBOA and MBOA decreased over time. Neither DIMBOA-Glc, HDMBOA-Glc nor DIMBOA showed statistically significant differences between Ct0861-inoculated and mock-treated plants. However, a significant reduction in the antimicrobial product MBOA was detected at 11 dpi in Ct0861-colonised roots (*P=*0.028). Overall, these results suggest a temporal regulation of BXs accumulation in the apoplast, with Ct0861 colonisation specifically affecting MBOA levels, with a transient decrease at the intermediate time point (11 dpi).

**Figure 4.**
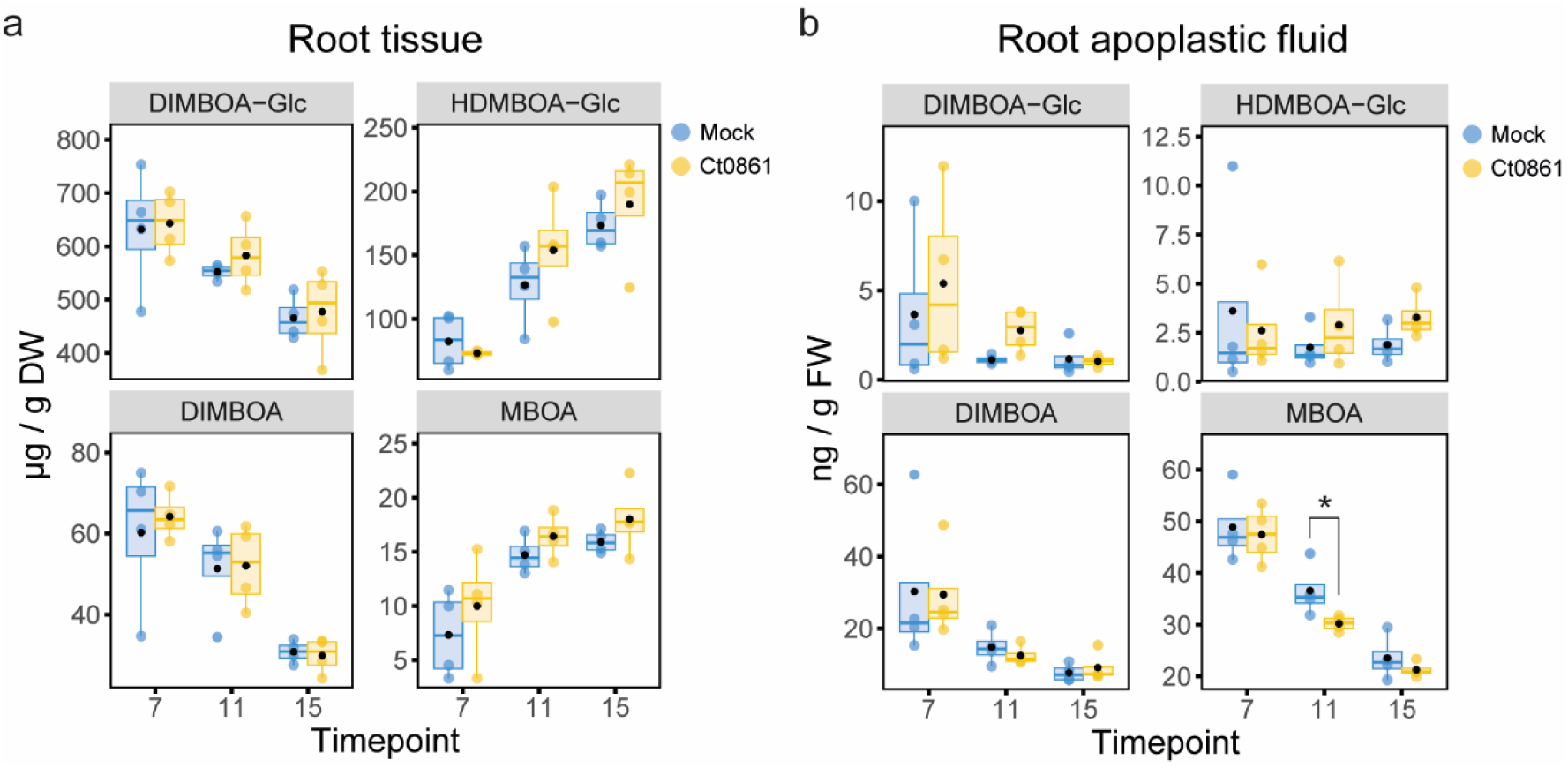
*Colletotrichum tofieldiae* strain Ct0861 alters 6-methoxy-benzoxazolin-2-one (MBOA) accumulation in maize (*Zea mays*) root apoplastic fluid. **a)** Concentrations of 2,4-dihydroxy-7-methoxy-1,4-benzoxazin-3-one glucoside (DIMBOA-Glc), 2-hydroxy-4,7-dimethoxy-1,4-benzoxazin-3-one glucoside (HDMBOA-Glc), 2,4-dihydroxy-7-methoxy-1,4-benzoxazin-3-one (DIMBOA) and MBOA in whole-root tissue from mock- and Ct0861-inoculated maize seedlings at 7, 11, and 15 days post-inoculation. Metabolite concentrations are expressed as µg/g dry weight (DW). **b)** Concentrations of the same benzoxazinoid-related metabolites in root apoplastic fluid collected from mock- and Ct0861-inoculated maize seedlings at 7, 11, and 15 dpi. Metabolite concentrations are expressed as ng/g fresh weight (FW). Internal line in box-plot boxes indicates the median or second quartile (Q2), upper line of the boxes indicates the third quartile (Q3) and lower line the first quartile (Q1) of the data. Mean is represented by a black dot. Asterisks denote statistically significant differences in Student’s t-test (*P* < 0.05).

### Ct0861 and CgM1.001 colonisation induce different tissue-specific defence transcriptional responses

To evaluate whether the plant responses observed during the early interaction with Ct0861 represent a general response to fungal colonisation, we inoculated maize seeds with spores of *C. graminicola* strain CgM1.001. *C. graminicola* is a fungal pathogen closely related to *C. tofieldiae* known to cause anthracnose stalk rot and leaf blight by colonising maize through leaves and roots (Sukno *et al*., 2008). Gene expression in roots and shoots was analysed at 7 dpi and compared with mock-treated plants (**Fig. S3**). Plants infected with CgM1.001 were smaller than mock or Ct0861 inoculated plants, with shorter roots and shoots, confirming the detrimental effect of this fungus on maize (**Fig. S3a**). In the roots, we observed a downregulation of *BX1* in plants treated with either Ct0861 or CgM1.001 (**Fig. S3b**), suggesting that this response is not specific to the interaction with Ct0861, but a broader response of a fungal inoculation. Conversely, genes involved in KXs, DXs and ZXs were upregulated, with a stronger expression increase in roots infected by CgM1.001. This differential gene expression correlates with the increased fungal biomass in CgM1.001-infected roots compared with fungal biomass amount in Ct-infected roots (**Fig. S3c**). In the shoots, KXs, DXs and ZXs genes were induced to a similar extent in response to both fungi (**Fig. S3b**). However, in contrast with the root, *BX1* was downregulated in the shoot only in CgM1.001-infected plants. Relative fungal biomass was also higher in shoots of CgM1.001-infected plants than in Ct0861-inoculated plants (**Fig. S3c**). Together, these results indicate that maize defence transcriptional responses to fungal colonisation are tissue-dependent and may reflect both the colonisation levels and the distinct interaction strategies of the beneficial endophyte Ct0861 and the pathogen CgM1.001.

### Ct0861 and CgM1.001 display distinct tolerance to MBOA

The metabolites produced by the benzoxazinoid pathway have been reported to be toxic to several phylogenetically distinct fungal species (Niculaes *et al*., 2018). We aimed to evaluate the inhibitory capacity of BX compounds on both Ct0861 and CgM1.001. To do this, we cultured the fungi on PDA plates amended with MBOA at different concentrations, observing a clear dose-dependent inhibitory effect (**Fig. 5a,b**). However, for CgM1.001, the inhibitory effect was observed only at 1 mg/mL, whereas growth was even enhanced at 0.25 mg/mL. Several maize-associated bacteria have been described to metabolise MBOA to 2-amino-7-methoxy-3H-phenoxazin-3-one (AMPO) resulting in orange pigmentation (Thoenen *et al*., 2024). In our experiments, the orange pigmentation was observed in CgM1.001 cultures grown on MBOA-amended media, but not in Ct0861 (**Fig. 5c**).

**Figure 5.**
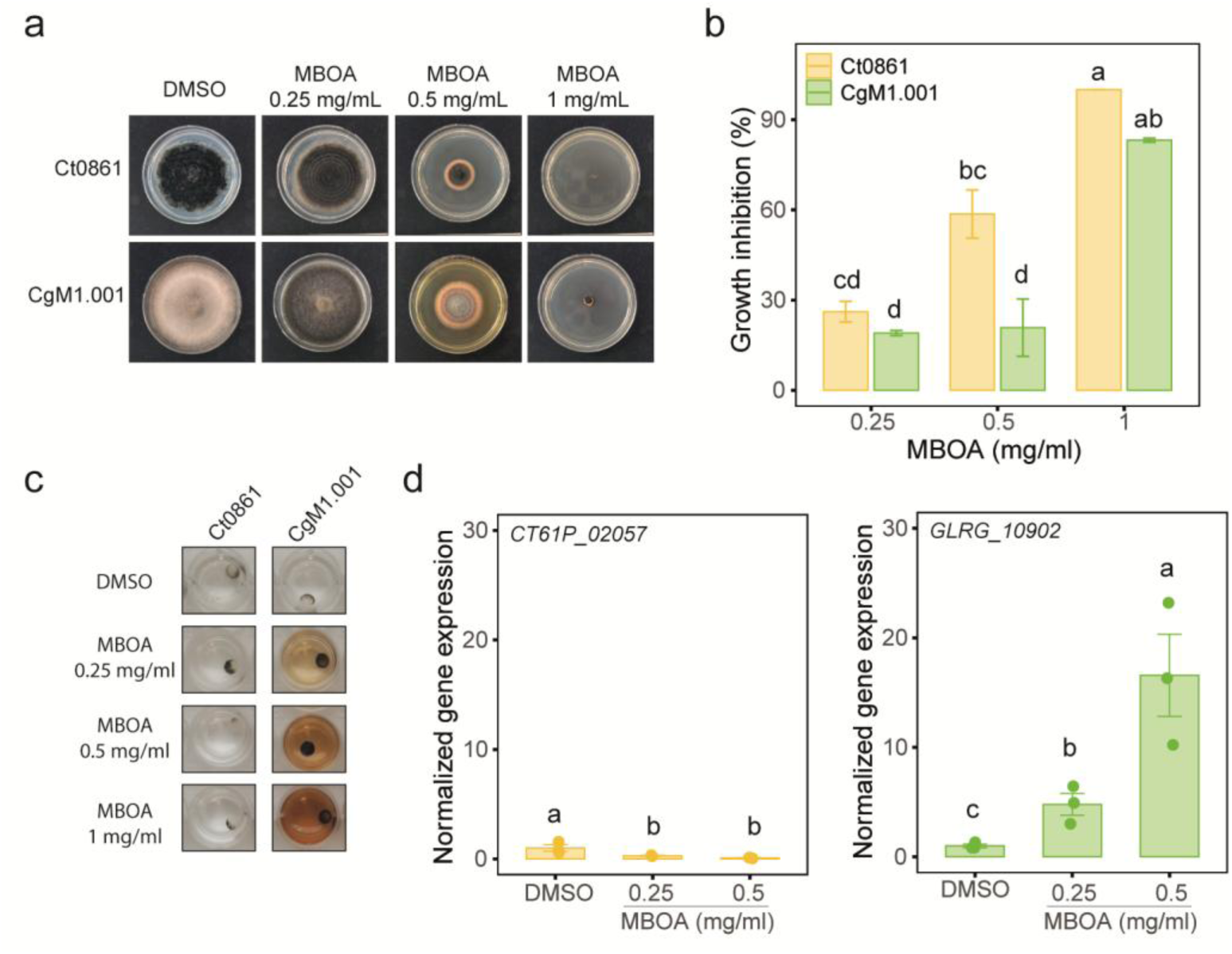
*Colletotrichum tofieldiae* strain Ct0861 and *Colletotrichum graminicola* strain CgM1.001 display distinct responses to methoxybenzoxazolin-2(3H)-one (MBOA) exposure. **a)** Representative images of Ct0861 and CgM1.001 colonies grown on PDA medium supplemented with dimethyl sulfoxide (DMSO) or increasing concentrations of MBOA. **b)** Percentage of fungal growth inhibition at the indicated MBOA concentrations relative to the corresponding DMSO control. Bars represent the mean ± standard deviation of 3 biological replicates. Different letters indicate statistically significant differences among fungus–MBOA treatment combinations, as determined by one-way ANOVA followed by Fisher’s least significant difference (LSD) test with Bonferroni correction (*P* < 0.05). **c)** Representative images of a liquid culture of Ct0861 and CgM1.001 grown supplemented with DMSO or increasing concentrations of MBOA. **d)** Relative expression levels of *CT61P_02057* in Ct0861 and *GLRG_10902* in CgM1.001 after exposure to the indicated MBOA concentrations, quantified by RT–qPCR. Bars represent the mean ± standard error. Different letters indicate statistically significant differences among fungus–MBOA treatment combinations, as determined by one-way ANOVA followed by LSD test with Bonferroni correction (*P* < 0.05).

Since *C. graminicola* has been reported to possess a putative xenobiotic-detoxification cluster homologous to the *Fusarium verticillioides FDB1* cluster (Glenn *et al*., 2016), we next examined whether this response was associated with the induction of a detoxification-related gene. The *C. graminicola* cluster includes *GLRG_10902*, a homolog of *F. verticillioides MBL1*, which encodes a metallo-β-lactamase required for the first step of BOA detoxification (Glenn *et al*., 2016). We identified *Ct61P_02057 as* the putative homolog of *GLRG_10902* in the Ct0861 genome and analysed the expression of both *GLRG_10902* and *Ct61P_02057* genes in fungal cultures grown in liquid medium supplemented with MBOA. RT-qPCR analysis showed that *GLRG_10902* was strongly induced by MBOA in a dose-dependent manner, with significantly higher expression at 0.25 and 0.5 mg/mL than in the DMSO control (**Fig. 5d**). In contrast, *Ct61P_02057* was not induced by MBOA and showed reduced expression in MBOA-treated samples (**Fig. 5d**). These results show that CgM1.001 and Ct0861 differ not only in their tolerance to MBOA, but also in the transcriptional response of candidate homologues associated with MBOA detoxification.

### Ct0861 colonisation and growth-promoting effects are enhanced in a BX-deficient maize mutant

Given the toxicity of BXs to Ct0861, we hypothesised that their absence in the plant would enhance Ct0861 colonisation and consequently affect the outcome of the interaction. To investigate this, we used the maize *bx1::DS* mutant and its near-isogenic W22 background as a control. The *bx1::DS* mutant lacks benzoxazinoid production due to a transposon insertion in the *BX1* gene, which disrupts the initial step of converting indole-1,3-glycerol phosphate to indole (Tzin *et al*., 2015; Hu *et al*., 2018).

Quantitative PCR analysis revealed a significant increase in Ct0861 biomass within the roots of the *bx1::DS* mutant compared to the wild-type W22 line plants grown *in vitro* (**Fig. 6a**). However, no visible disease symptoms were observed, even after prolonged growth of the inoculated plants in the greenhouse for one month (**Fig. 6b**). Interestingly, inoculation with Ct0861 resulted instead in enhanced plant growth promotion in the *bx1::DS* mutant (**Fig. 6c**). Indeed, Ct0861 inoculation of *bx1::DS* plants increased root length, root fresh weight, shoot length and shoot fresh weight by 12.23%, 14.12%, 26.7%, and 28.68%, respectively, compared with mock controls. All increases were statistically significant except for root length (**Fig. 6c**). Moreover, the increases in root length, root fresh weight and shoot fresh weight were significantly greater than those observed in the Ct0861 inoculated wild type W22 plants which exhibited increments of 4.92 %, −1.09%, 26.83% and 13.93% in root length, root fresh weight, shoot length and shoot fresh weight, respectively. Collectively, these results indicate that BX deficiency amplifies the plant growth-promoting activity of Ct0861 without triggering disease symptoms. In contrast, seed inoculation of *bx1::DS* with pathogenic CgM1.001 did not increase fungal colonisation relative to W22 (**Fig. S4a**), nor did it exacerbate its detrimental effect on the plant (**Fig. S4b**), supporting the idea that BXs are not the main maize defences acting against this pathogen.

**Figure 6.**
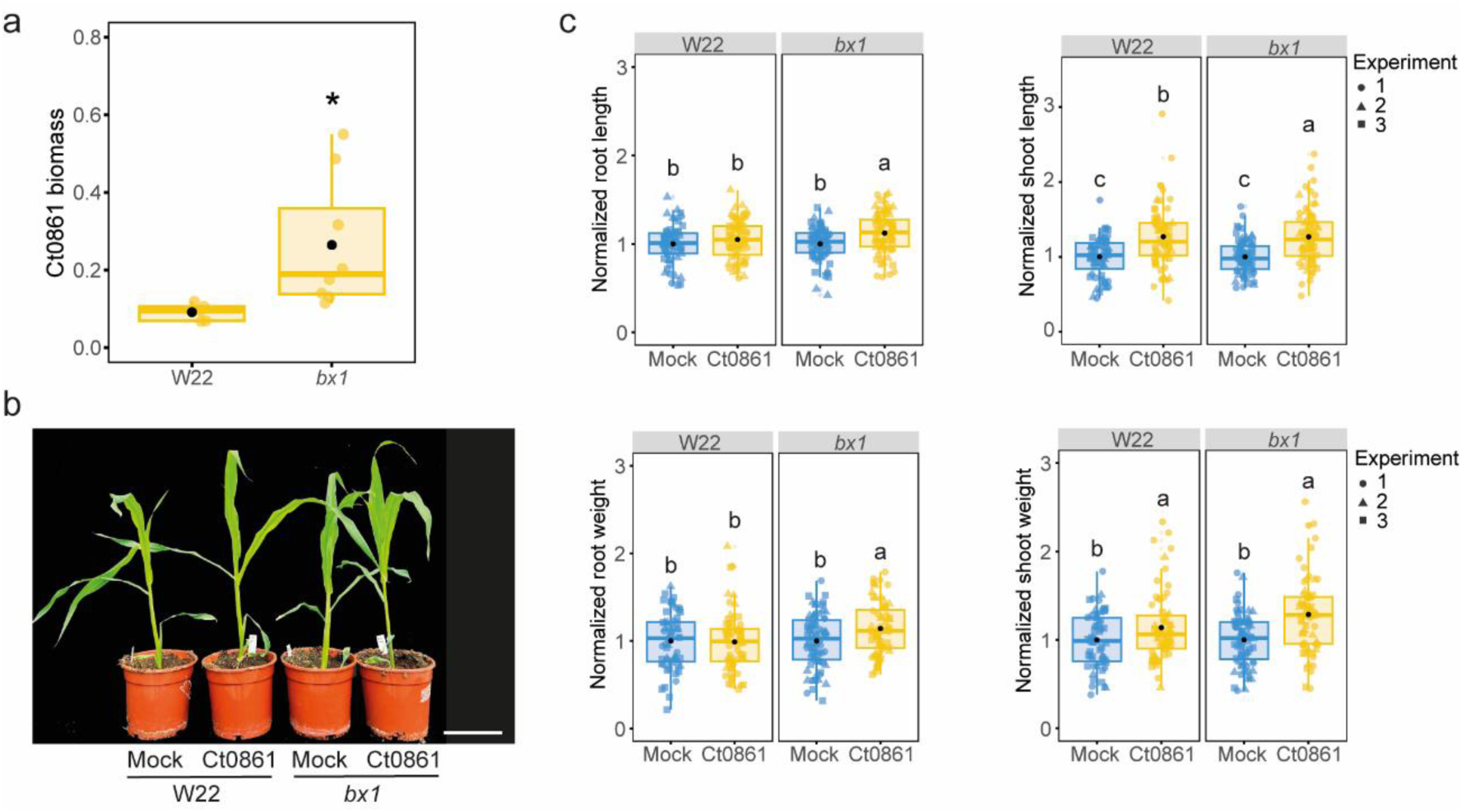
*Colletotrichum tofieldiae* strain Ct0861 shows enhanced colonisation and plant growth-promoting activity in a benzoxazinoid-deficient mutant. **a)** Ct0861 biomass in roots of the maize (*Zea mays*) wild-type line W22 and the *bx1::DS* mutant at 7 days post-inoculation (dpi) of plants grown *in vitro*. Fungal biomass was quantified by qPCR using Ct0861-specific primers and normalised to the maize *BETA-TUBULIN* gene. The asterisk indicates a statistically significant difference between genotypes (Student’s t-test, *P* < 0.05). **b)** Representative images of mock-treated and Ct0861-inoculated W22 and *bx1::DS* maize plants at 30 dpi. Scale bar = 18 cm. **c)** Root length, shoot length, root weight, and shoot weight of mock-treated and Ct0861-inoculated W22 and *bx1::DS* maize plants grown *in vitro* at 7 dpi. Values were normalised to the mean of the corresponding mock-treated genotype. Internal line in box-plot boxes indicates the median or second quartile (Q2), upper line of the boxes indicates the third quartile (Q3) and lower line the first quartile (Q1) of the data. Mean is represented by a black dot. Different letters indicate statistically significant differences among treatments, as determined by one-way ANOVA followed by LSD test with Bonferroni correction (*P* < 0.05).

## DISCUSSION

### Maize responses to beneficial Ct0861 colonisation are strongly tissue- and time-dependent

In this study, we show that the initial steps of maize colonisation by the beneficial fungal endophyte Ct0861 are accompanied by a dynamic reprogramming of the host transcriptome. By combining transcriptomic analyses across two tissues and two stages of colonisation, our study provides a spatiotemporal view of the early maize responses to this beneficial fungus. This approach reveals that host reprogramming is not uniform across the plant but instead is strongly tissue specific and dynamic over time, with roots showing the most pronounced response at 7 dpi (**Fig. 1b,d**), coinciding with increased fungal biomass (**Fig. 1c**). In this respect, our work extends previous transcriptomic studies of beneficial fungal interactions in maize, which have largely focused on single tissues (predominantly roots) and narrower time windows (Malinich et al., 2019; Sun et al., 2023; Zhang et al., 2025; Zhang et al., 2018).

### Ct0861 colonisation is associated with reprogramming of maize defence metabolism

One of the most prominent responses to Ct0861 colonisation is the modulation of the expression of biosynthetic genes involved in maize defence metabolism. Ct0861-colonised plants showed the repression of BX-related genes alongside the concurrent induction of genes associated with terpenoid pathways, including KXs, ZXs and DXs (**Fig. 2a,c**). This pattern is consistent with the highly modular organisation of maize biochemical immunity, where BXs serve as the primary constitutive chemical defence in seedlings. The decline of BXs as the plant matures facilitates a temporal transition to a diverse array of specialised inducible defensive metabolites, including terpenoids, which dominate at later developmental stages (Yasmin *et al*., 2024; Zhou *et al*., 2024). Thus, colonisation by Ct0861 appears to trigger an early onset of this temporal transition of defensive molecules, accelerating the deployment of specialised terpenoid defences over the seedling-specific BX-based defence layer.

A similar reconfiguration of maize defence metabolism has previously been described following plant treatments with fungal elicitors, suggesting that the host response to Ct0861 reflects part of a general defence reprogramming against fungi. Ahmad et al. (2011) showed the downregulation of BX biosynthetic genes in leaves upon elicitation with the fungal cell wall component chitosan. Similarly, Ding et al. (2020) observed a dramatic suppression of early BX biosynthetic steps alongside a strong concurrent induction of acidic terpenoid pathways (including KXs and ZXs) following elicitation with heat-killed *Fusarium venenatum* hyphae. Ding et al. (2020) suggested that this response reflects a metabolic *trade-off* in which resources are reallocated from constitutive, indole-derived defences toward the deployment of inducible, highly potent terpenoid antibiotic cocktails that optimise defence effectiveness. This interpretation is further supported by evidence of functional specialisation and genetic pleiotropy in maize antifungal defences. Yang et al. (2019) found that resistance to *Exserohilum turcicum* mediated by ZmWAK-RLK1 correlates with reduced BXs content, suggesting that while BXs are critical for insect defence, they may be ineffective or even counterproductive against certain fungal pathogens compared to more specialised terpenoid responses. Indeed, many described maize defences against fungal pathogens rely on terpenoid phytoalexins rather than benzoxazinoids, likely because several maize-associated fungi show tolerance to benzoxazinoids (Glenn *et al*., 2001; Saunders & Kohn, 2009; Christensen *et al*., 2018; Saldivar *et al*., 2023).

### Specific defence responses differentiate beneficial Ct0861 from pathogenic CgM1.001

A similar repression of *BX1* alongside the induction of terpenoid-associated genes was also detected in response to the hemibiotrophic pathogen *C. graminicola* strain CgM1.001 (**Fig. S3b**). This is consistent with previous reports of reduced *BX1* expression during the early stages of *C. graminicola* infection (Balmer *et al*., 2013), and supports the idea that this transcriptional shift forms part of a broader host response to fungal colonisation rather than a hallmark unique to Ct0861. Nonetheless, our data reveal both quantitative and qualitative differences between Ct0861 and CgM1.001, with the pathogenic interaction displaying stronger transcriptional activation, higher fungal biomass and associated disease symptoms (**Fig. S3**). Moreover, while *BX1* repression occurred in roots in response to both fungi, shoot repression was only detected in plants colonised by CgM1.001 (**Fig. S3b**). *C. graminicola* is generally considered a leaf and stalk pathogen, although it can infect roots and subsequently cause systemic infection of aerial tissue (Sukno *et al*., 2008). In contrast, Ct0861 predominantly colonises roots, only occasionally invading *A. thaliana* shoots systemically (Hiruma *et al*., 2016; Díaz-González *et al*., 2020). Thus, while both *BX* repression and the transcriptional activation of terpenoid pathways appear proportional to the extent of fungal colonisation, terpenoid defences are systemically elicited, whereas BX repression reflects a localised response to fungal presence within the tissue.

The outcome of the interaction depends not only on host transcriptional reprogramming but also on fungal traits, such as tolerance to or detoxification of host-derived metabolites. Glenn et al. (2016) provided evidence for the horizontal transfer of the *FDB1* cluster required to detoxify BXs from *Fusarium* to *C. graminicola*, although they did not test its functional activity in the latter species. Our analyses confirm the ability of CgM1.001 to tolerate MBOA and suggest its conversion to AMPO (**Fig. 5**). This tolerance is linked to the upregulation of *GLRG_10902*, a homologue of the *F. verticillioides* MBL1 metallo-β-lactamase present in the *FDB1* cluster and required for the initial step of BOA detoxification (Glenn *et al*., 2016). Although we identified a homologue of *GLRG_10902* in the Ct0861 genome, this gene was not upregulated in response to MBOA, nor was any detoxification capacity observed for Ct0861 (**Fig. 5**). Hence, while part of the *FDB1* cluster may be conserved in Ct0861, its function in BX detoxification has not been retained. Indeed, a clear correlation exists between host BX exposure and the fungal capacity to detoxify these compounds, and it is assumed that BX tolerance provides a selective advantage for grass-specialised pathogens (Saunders & Kohn, 2009; Kettle *et al*., 2015; Glenn *et al*., 2016). Although *C. graminicola* and *C. tofieldiae* are closely related species (Hacquard *et al*., 2016), *C. graminicola* is a specialised maize pathogen (Crouch *et al*., 2006), whereas *C. tofieldiae* is a generalist endophyte capable of colonising a diverse array of dicot and monocot hosts (Díaz-González *et al*., 2020), and may not have been subjected to the selective pressure required to maintain BX detoxification mechanisms.

### Indole-derived defences against Ct0861 are subject to host-species-specific regulation

The repression of BX biosynthesis in Ct0861 colonised roots contrasts with previous findings in *A. thaliana*, where Trp-derived IGs are induced in response to Ct0861 (Hiruma *et al*., 2016; Hacquard *et al*., 2016). Although maize BXs and *A. thaliana* IGs belong to distinct, lineage-specific, defence systems, both represent major indole-derived chemical interfaces through which plants modulate fungal colonisers (Erb & Kliebenstein, 2020; Schlaeppi *et al*., 2021). In *A. thaliana*, disruption of Trp-derived secondary metabolism, and in particular IG-associated pathways, compromises host control over Ct0861 growth, shifting the interaction towards a detrimental outcome (Hiruma *et al*., 2016). Our results therefore point to a marked functional divergence in indole-mediated defences between these two host species.

Since indole is a shared precursor, it is tempting to hypothesise that the modulation of indole-related defences could influence auxin-related metabolism. In *A. thaliana*, Frerigmann et al. (2021) proposed that Ct0861 might utilise IG hydrolysis to generate indole-3-acetonitrile (IAN) and IAA via NIT1–3 nitrilase activity. However, *nit1/2/3* and *NIT2* RNAi mutants displayed no defect in Ct0861-induced growth promotion, demonstrating that this metabolic connection is dispensable for the beneficial activity of the fungus. Thus, because Ct0861 restores auxin signalling under phosphate-starved conditions, the authors concluded that either an alternative metabolic link exists between IG hydrolysis products and auxin biosynthesis, or this effect occurs independently of IG metabolism (Frerigmann *et al*., 2021). In analogy, *BX* repression does not appear to reflect a rerouting of indole metabolism, as we detected neither the transcriptional activation of alternative indole-producing branches (**Fig. 3a**) nor changes in IAA accumulation (**Fig. 3b**). Thus, interconnections between indole-derived defences and IAA do not appear to underlie the growth-promotion effect in either host species.

### Reduced benzoxazinoid defences favour beneficial Ct0861 colonisation in maize

The downregulation of BX biosynthetic genes upon Ct0861 colonisation was followed by a reduction in MBOA levels within the root apoplast (**Fig. 4b**), which could facilitate endophytic root colonisation by Ct0861. Consistent with this hypothesis, disruption of BX biosynthesis in the *bx1::DS* mutant led to increased fungal biomass (**Fig. 6a**). However, in contrast to findings in *A. thaliana* (Hiruma *et al*., 2016), far from being detrimental, this increase in fungal biomass was accompanied by an enhanced growth-promotion phenotype (**Fig. 6b,c**). Also contrary to expectations, CgM1.001 showed neither an enhanced fungal biomass accumulation nor any detrimental effects during its interaction with the *bx1::DS* mutant (**Fig. S4**), indicating that BXs do not play a major role in maize defences against this pathogen. Taken together, these results support a model in which the partial attenuation of BXs in response to Ct0861 favours fungal proliferation and confers physiological benefits to maize, without increasing susceptibility to specialised pathogens.

In summary, the positive interaction between Ct0861 and maize may benefit from a general host response to fungi, consisting of the downregulation of BX biosynthesis alongside the concurrent shift towards terpenoid-mediated defences. This phenomenon may have significant consequences for the interaction of maize seedlings with beneficial fungi, opening a temporal window of reduced BX levels that facilitates microbial establishment within the root system. This mechanism would be particularly relevant for generalist beneficial microorganisms lacking tolerance to BXs, especially given that these metabolites do not appear to play a key role in restricting specialised pathogens. These interpretations align with the broader view of BXs as multifunctional maize metabolites that not only function in direct defence against antagonists but also contribute to structuring plant-associated microbial communities and belowground ecological interactions (Kudjordjie *et al*., 2019; Cadot *et al*., 2021; Gfeller *et al*., 2023). Beyond acting as a strict on/off switch for compatibility, BXs regulation may fine-tune the extent of fungal accommodation and, consequently, the magnitude of the resulting beneficial effects.

## Supporting information

Table S1. Oligonucleotide primers

Table S2. Differentially expressed genes

Table S3. Functional enrichment analysis

## ACKNOWLEDGEMENTS

We thank Dr. Georg Jander (Boyce Thompson Institute, NY, USA), Dr. Monika Frey (Technical University of Munich, Germany), Dr. Michaela Matthes (University of Bonn, Germany), Dr. Christelle Robert (University of Bern, Switzerland) and Dr. Ana Butrón (MBG-CSIC, Spain) for generously providing maize seeds (mutant and wild-type lines). We also thank Dr. Christelle Robert and Dr. Jurriaan Ton (University of Sheffield University, UK) for providing BX standards, as well as to Dr. Christelle Robert for her advice on BX metabolomic quantification. We are grateful to Dr. Ana Butrón for her valuable advice on maize cultivation and handling, and to Dr. Serenella Sukno (University of Salamanca, Spain) for providing CgM1.001 strain.

The authors acknowledge the use of artificial intelligence tools (Microsoft Copilot, ChatGPT, Google Gemini, Consensus and Claude), for language editing and proofreading. All AI-assisted text was critically reviewed and revised by the authors to ensure accuracy, clarity, and scientific integrity.

This research was supported by grants PID2021-123697OB-I00 and PID2024-161830OB-I00 (PI S.S.) funded by MCIN/AEI/10.13039/501100011033/ and ERDF/EU. C.G.-S. was supported by Severo Ochoa Program for Centres of Excellence at CBGP (CEX2020-000999-S-20-4) and grant PRE2022-103983 from MCIN/AEI/10.13039/501100011033 and ESF+. B.W. was supported by a predoctoral fellowship of the China Scholarship Council. L.R.-C. was supported by the Severo Ochoa Program for Centres of Excellence at CBGP (CEX2020-000999-S-20-4) funded by MCIN/AEI/ 10.13039/501100011033. Technical assistance by Sylwia Szygut was funded by grant PEJ-2024-TL_BIO-33107 from the Community of Madrid and ESF+.

## AUTHOR CONTRIBUTIONS

C.G.S., S.D.G. and S.S. conceived and designed the study. C.G.S., S.D.G. and B.W. performed the experiments and collected the data. C.G.S., S.D.G. and B.W. analysed and interpreted the data. S.G.B. and L.R.C. provided bioinformatic support. V.F. conducted metabolomic analyses. C.G.S., S.D.G. and S.S. wrote the original draft of the manuscript. S.S. supervised the research. All authors reviewed and edited the manuscript and approved the final version.

## COMPETING INTERESTS

S.S. is an inventor on patent ES 2439393 and its associated patent family, assigned to UPM and PlantResponse Biotech S.L. C.G.-S., S.D.-G., and S.S. are inventors on patent ES 2969994 and its associated patent family, assigned to UPM and Plant Response Inc. No funding from Plant Response Biotech S.L. or Plant Response Inc. was received for this research, and these companies had no involvement in the writing of this manuscript or the decision to submit it for publication. W.B. S. G.-B., V.F. and L. R.-C. declare no competing interests.

## DATA AVAILABILITY STATEMENT

The raw RNA-seq data generated in this study have been deposited in the NCBI Sequence Read Archive (SRA) under BioProject accession PRJNA1504184. Differential expression and functional enrichment datasets are provided in Supporting Information Tables S2 and S3.

