## Supporting Information for "Reduced benzoxazinoid defences favour maize beneficial colonisation by *Colletotrichum tofieldiae*"

### Index

- Supplementary Figures
- Supplementary Tables

### Supplementary Figures

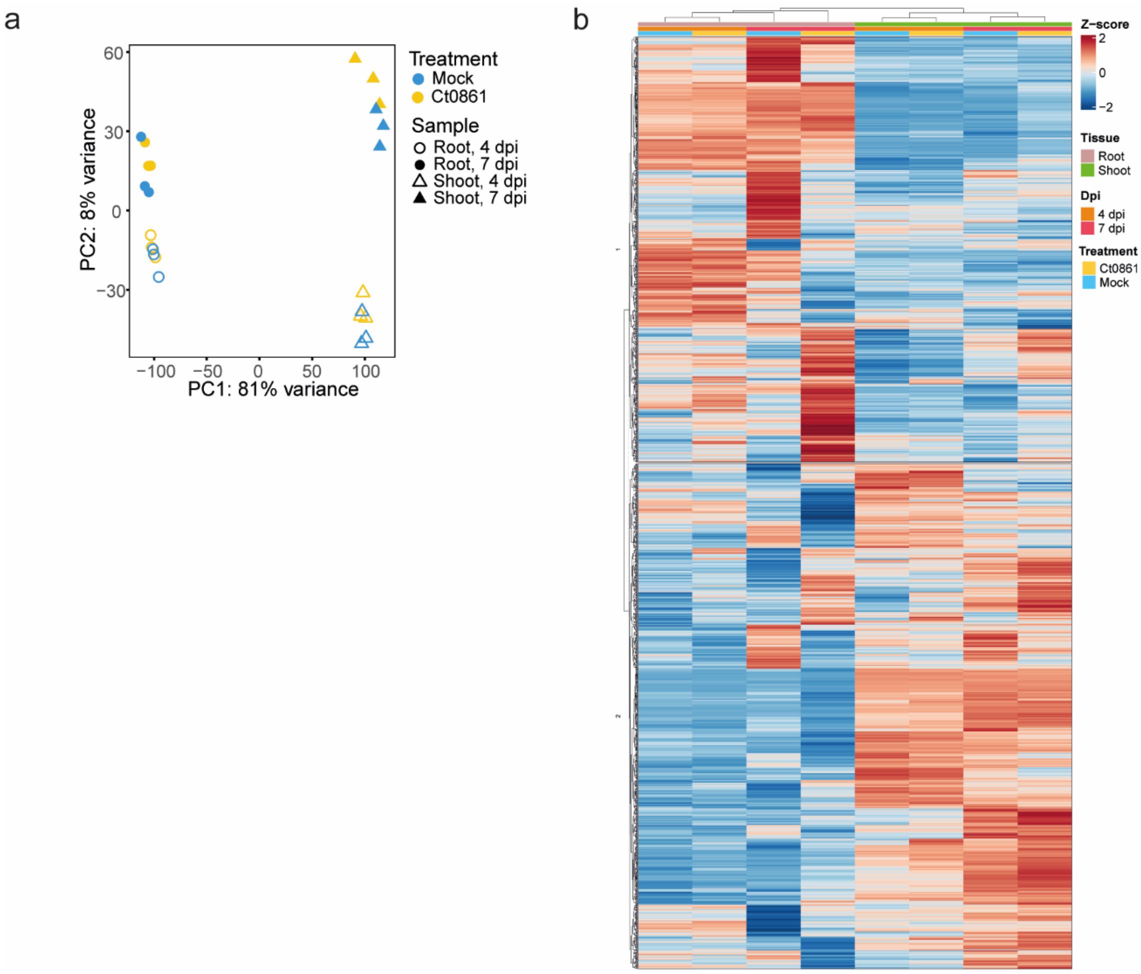


**Figure S1. Global transcriptomic profiles of mock-treated and *Colletotrichum tofieldiae* strain Ct0861-inoculated maize (*Zea mays*) roots and shoots. a)** Principal component analysis of RNA-seq data from roots and shoots of mock-treated and Ct0861-inoculated maize seedlings at 4 and 7 days post-inoculation (dpi). Colours indicate treatment, circles and triangles indicate roots and shoots, respectively, and open and filled symbols indicate samples collected at 4 and 7 dpi, respectively. The percentages shown on the axes indicate the proportion of total variance explained by each principal component. **b)** Heat map showing the expression profiles of differentially expressed genes identified across all comparisons between Ct0861-inoculated and mock-treated maize roots and shoots at 4 and 7 dpi. Normalised expression values were averaged by condition and transformed into row-scaled Z-scores. Genes and samples were hierarchically clustered using Euclidean distance.

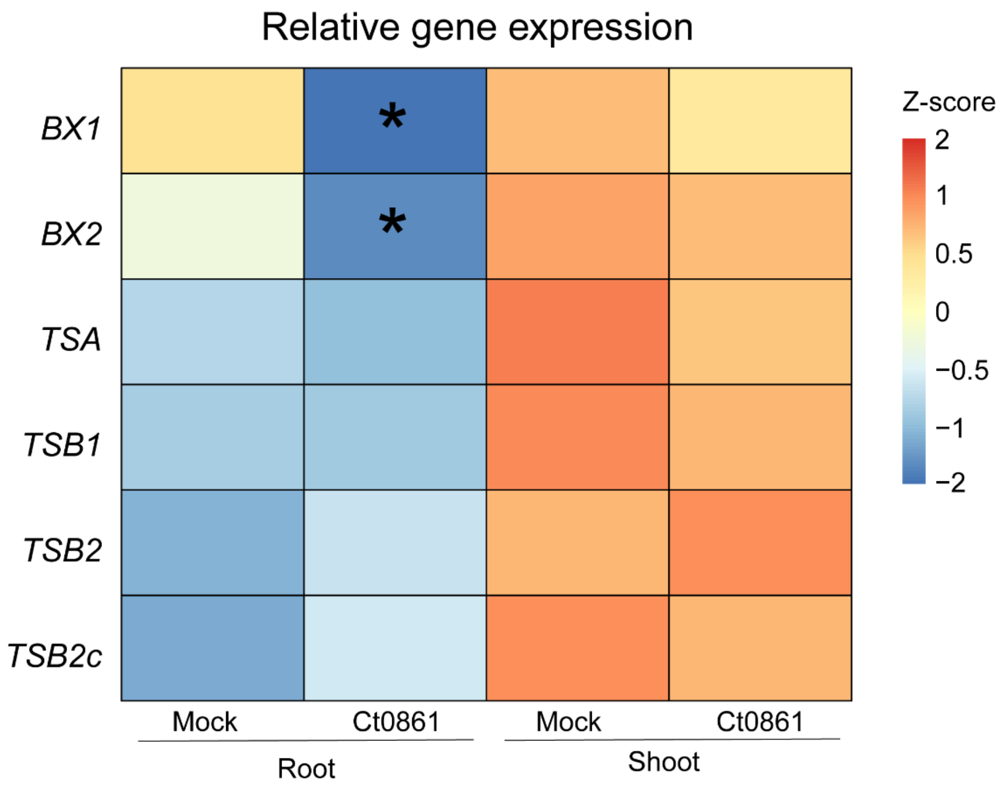


**Figure S2. *Colletotrichum tofieldiae* strain Ct0861 alters benzoxazinoid-related gene expression without affecting the tryptophan biosynthesis pathway in maize (*Zea mays*).** Heat map showing the relative expression of *BX1*, *BX2*, *TSA*, *TSB1*, *TSB2*, and *TSB2c* in roots and shoots of mock-treated and Ct0861-inoculated maize seedlings at 7 days post-inoculation (dpi). Gene expression was quantified by RT–qPCR using gene-specific primers and normalised to *ACTIN*. Values are displayed as row-scaled Z-scores calculated from the relative expression values of each gene. Asterisks indicate statistically significant differences between Ct0861-inoculated and mock-treated samples within the corresponding tissue, as determined by Student's t-test ($P < 0.05$).

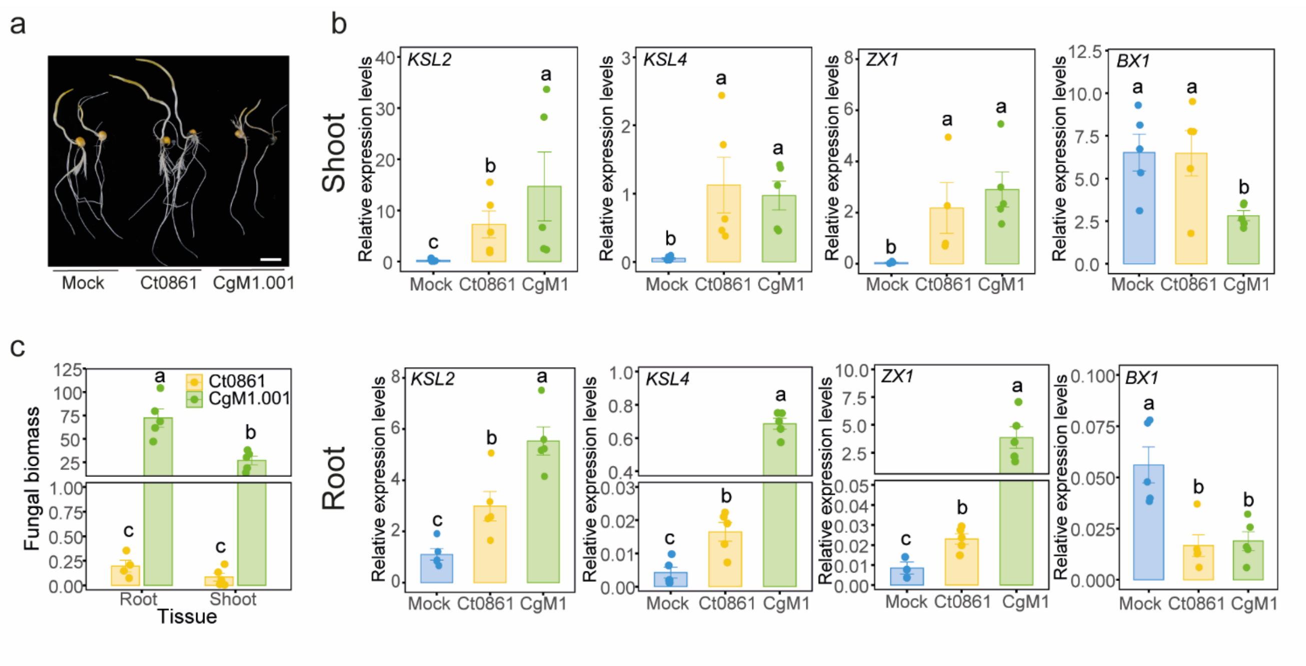


**Figure S3. *Colletotrichum tofieldiae* strain Ct0861 and *Colletotrichum graminicola* strain CgM1.001 differentially colonise maize (*Zea mays*) tissues and modulate specialised defence-related genes**. **a)** Representative images of maize seedlings inoculated with *Ct0861*, *CgM1.001*, or mock-treated at 7 days post-inoculation (dpi). Scale bar = 2 cm. **b)** Relative expression levels of *KSL2*, *KSL4*, *ZX1* and *BX1* in shoots (top) and roots (bottom) of mock-, *Ct0861*-, and *CgM1.001*-inoculated maize seedlings. Gene expression was quantified by RT–qPCR using gene-specific primers and normalised to *ZmACTIN*. Bars represent the mean ± standard error. Different letters indicate statistically significant differences determined by one-way ANOVA followed by LSD test with Bonferroni correction ($P < 0.05$). **c)** Quantification of fungal biomass in roots and shoots of maize seedlings inoculated with *Ct0861* or *CgM1.001*. Fungal biomass was determined by qPCR using fungus-specific primers and normalised to the maize β-tubulin gene. Bars represent the mean ± standard error.

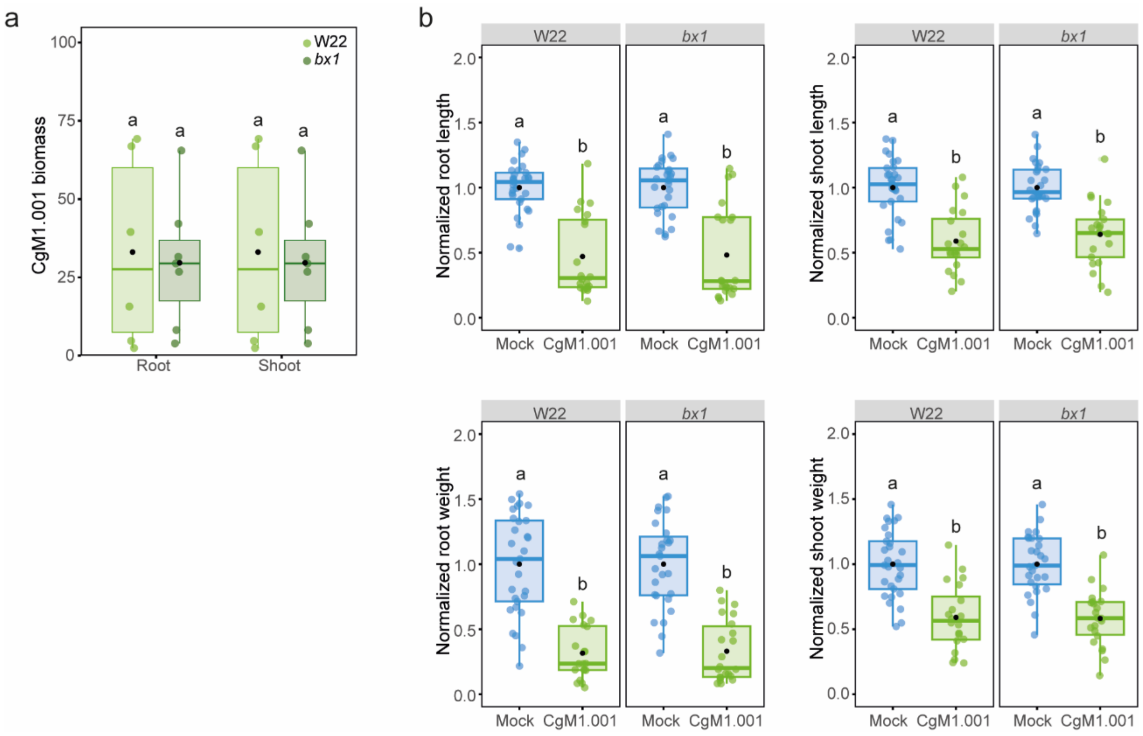


**Figure S4. Colonisation and growth effects of *Colletotrichum graminicola* CgM1.001 in W22 and *bx1* maize (*Zea mays*) plants. a)** CgM1.001 biomass in roots of the maize wild-type line W22 and the *bx1* mutant at 7 days post-inoculation (dpi). Fungal biomass was quantified by qPCR using CGM1.001-specific primers and normalised to the maize β-tubulin gene. **b)** Root length, shoot length, root weight, and shoot weight of mock-treated and CgM1.001-inoculated W22 and *bx1* maize plants. Values were normalised to the mean of the corresponding mock-treated genotype. Internal line in box-plot boxes indicates the median or second quartile (Q2), upper line of the boxes indicates the third quartile (Q3) and lower line the first quartile (Q1) of the data. Mean is represented by a black dot. Different letters indicate statistically significant differences among treatments, as determined by one-way ANOVA followed by LSD test with Bonferroni correction ($P < 0.05$).

### Supplementary Tables

**Table S1. Oligonucleotide primers used in this study.** Primer names and nucleotide sequences (5′–3′), target genes, gene identifiers, species, and source references are indicated. Primers without a previous reference were designed in this study.

**Table S2. Differentially expressed genes identified in each treatment compared with the corresponding mock control.** Differentially expressed genes (DEGs) were identified using DESeq2 and defined as genes with an absolute log2 fold change ($|log2FC|$) ≥ 1 and an adjusted *P* value < 0.05. Results for each comparison are provided in separate worksheets.

**Table S3. Functional enrichment analysis of differentially expressed genes for each comparison.** Gene Ontology (GO) terms and Kyoto Encyclopedia of Genes and Genomes (KEGG) pathways significantly enriched among differentially expressed genes are reported separately for each treatment compared with the corresponding mock control. Significantly enriched categories were defined using an adjusted *P* value < 0.05. GO terms are classified according to the three GO domains: biological process (BP), molecular function (MF), and cellular component (CC). Results for each comparison are provided in separate worksheets.
